# Reducing disease and producing food: Effects of 13 agrochemicals on snail biomass and human schistosomes

**DOI:** 10.1101/2021.01.05.425422

**Authors:** Christopher J E Haggerty, Neal Halstead, David J. Civitello, Jason R Rohr

## Abstract

Agrochemical use is predicted to increase 2-5 fold by 2050 to meet food demand. Evidence suggests that agrochemical pollution could increase snails that transmit the disease schistosomiasis to 250 million people, but most agrochemicals remain unexamined. Here we quantify the relative effects of fertilizer, six insecticides, and six herbicides on snail genera responsible for 90% of global schistosomiasis cases. We identified fertilizers and 4 of 6 insecticides as high risk for increasing snail biomass by increasing snail resources (vegetative habitat and periphytic food) and reducing snail predators, respectively. Herbicides generally had negative effects on snails by reducing vegetative habitat, with two herbicides increasing snails in the absence of aquatic vegetation. Parasite production, which reflects human infection risk, scaled positively to snail biomass. Our findings suggest that fertilizers and insecticides are more likely to increase human schistosomiasis than herbicides and revealed several low risk agrochemicals that might increase crop production without increasing schistosomiasis.

---

Recent evidence suggests that increasing agriculture to meet the growing global food demand could simultaneously increase human infectious diseases (Rohr et al., 2019), necessitating studies that identify means of sustainably improving food production without increasing disease. One such disease that is positively associated with agricultural expansion and intensification is schistosomiasis. Schistosomiasis is a disease of humans caused by parasites released from freshwater snails, with an estimated 779 million people at risk globally (Vos et al., 2016). Second only to malaria in socioeconomic impact in tropical countries, schistosomiasis remains a neglected tropical disease in terms of research funding (May, 2007; Moran et al., 2009). Over 90% of global cases (∼200 million cases) occur in sub-Saharan Africa and involve *Biomphalaria* and *Bulinus* snail species as intermediate hosts (Gryseels, Polman, Clerinx, & Kestens, 2006; Steinmann, Keiser, Bos, Tanner, & Utzinger, 2006). Increased prevalence and geographic expansion of schistosomiasis in recent decades has been linked to expanding snail habitat and loss of native snail predators (Huang & Manderson, 1992; Savaya Alkalay et al., 2014; Steinmann et al., 2006). Infected snail hosts are key to disease transmission because each snail can release thousands of cercariae, the human-infectious life stage, daily. Parasite production by snails can be influenced by local aquatic factors (Haggerty, Bakhoum, Civitello, et al., 2020), and thus, human infection rates respond rapidly to changes in local ecological factors (Huang & Manderson, 1992), emphasizing the importance of freshwater ecology in infection dynamics.

Recent experimental evidence suggests that agrochemical pollution has the potential to influence several ecological factors in freshwater that directly or indirectly influence snail hosts that transmit human schistosomes. Both fertilizers and herbicides can increase periphyton resources that are needed for snail growth and cercarial production (Halstead et al., 2014), and have been linked to increased trematode infection intensities in frogs in agricultural wetlands (Rohr et al., 2008). Herbicides kill phytoplankton that absorb light from the water column, which can indirectly increase growth of periphyton, attached algae that is a major resource for snails (Halstead et al., 2018). Additionally, fertilizers and herbicides can increase and decrease, respectively, submerged vegetation that acts as primary habitat for snails and may influence snail predation. Invertebrate predators that consume snails and help regulate snail populations can be killed by some insecticides even at environmentally relevant concentrations, leading to top-down insecticide effects that increase snails (Halstead et al., 2014). Given that eliminating snail intermediate hosts is extremely difficult (Gryseels et al., 2006; Steinmann et al., 2006), reductions in snails and their cercariae, either by improving snail predator populations (Hofkin, Hofinger, Koech, & Loker, 1992; Mkoji et al., 1999; Savaya Alkalay et al., 2014; Sokolow, Lafferty, & Kuris, 2014) or limiting snail habitat, may help reduce schistosome transmission to humans and therefore, sustainably facilitate agricultural expansion and/or intensification. Understanding the strength of various agrochemicals on snails might be key to understanding recent increases in human schistosomiasis coincident with agricultural expansion and preventing any future surges (Hoover et al., 2020).

Agricultural expansion is predicted to increase agrochemical use 2-5 fold over the next few decades (Foley et al., 2011; Tilman, Balzer, Hill, & Befort, 2011; Tilman et al., 2001), yet laboratory microcosm studies of pesticides using species in isolation could miss the net effects of pesticides in natural communities. Community-level responses to contaminants can be mediated by interactions with other members of the community (Clements & Rohr, 2009; Rohr & Crumrine, 2005; Rohr, Kerby, & Sih, 2006). For example, submerged aquatic vegetation that acts as snail habitat might decrease the toxicity of certain insecticides by changing pH (Brogan & Relyea, 2014). Likewise, the strength of top-down regulation can be mediated by the presence of submerged aquatic vegetation, which can either decrease rates of predation by providing refugia for prey or increase access of predators to prey (Davis, Purrenhage, & Boone, 2012). Thus, because agrochemicals in natural environments are differently subject to the above influences, simulating natural communities using mesocosm experiments is needed to improve predictions of the direct and indirect effects of agrochemicals on freshwater communities, such as those containing snail hosts (Rohr, Salice, & Nisbet, 2016).

We created experimental natural freshwater pond communities to determine the predictability of top-down effects of insecticides and bottom-up effects of fertilizer or herbicides on the biomass of two genera of snail hosts that can transmit schistosomiasis (*Biomphalaria glabrata* and *Bulinus truncatus*). More specifically, we examined the top-down effects of six insecticides belonging to two insecticide classes (the organophosphates chlorpyrifos, malathion, and terbufos, and the pyrethroids esfenvalerate, λ-cyhalothrin, and permethrin), and the bottom-up effects of both fertilizer and six herbicides belonging to two herbicide classes (the chloroacetanilides acetochlor, alachlor, and metachlor, and the triazines atrazine, propazine, and simazine) in the presence of snail predators. We then performed a follow-up experiment using all six herbicides but without snail predators to determine if herbicide bottom-up effects on snails are only evident in the absence of strong top-down effects. In the follow-up experiment, we also examined if total parasite production by snails was predicted by their total biomass, and, thus, whether snail biomass changes associated with different agrochemical exposures could be related to human exposure to parasites.

## Results

### Experiment One

Pesticide type explained 43 percent of the variation in *Bi. glabrata* biomass (Table S4), whereas most of the variation in *Bu. truncatus* biomass (35 percent) was explained by pesticide identity nested within class nested within type (Table S4). Likelihood ratio tests determined that the above pesticide effects were significant for both snail species (Table S5). Model selection favored using pesticide type and pesticide nested within class within type for *Bi. glabrata* and *Bu. truncatus* biomass, respectively, and the lowest AICc values included a term for zero-inflation and treating water and solvent as a single control group for both snail species (Tables S6-S7). Model selection (Table S6) indicated significant positive fertilizer effects (bottom-up) on *Bi. glabrata* biomass (Fig. 1a; Table 1). In contrast, bottom-up effects of herbicides for *Bi. glabrata* biomass were generally not significant in the presence of predators (Table 1; *p* > 0.05). Insecticides were associated with positive top-down effects on *Bi. glabrata* biomass (Fig. 1a; Table 1). Similar to *Bi. glabrata*, fertilizers had significant positive bottom-up effects for *Bu. truncatus* biomass, after model selection (Table S7), while none of the six herbicides were significantly associated with *Bu. truncatus* biomass in the presence of predators (Fig. 1b; Table 1). Similar to *Bi. glabrata*, insecticides were associated with positive top-down effects on *Bu. truncatus* biomass (Fig. 1b). Four insecticides led to significant increases in average *Bu. truncatus* biomass relative to the controls (chlorpyrifos: 55% increase, terbufos: 36%, esfenvalerate: 28%, and permethrin: 61%), while one insecticide from each insecticide class had no significant effects (malathion and λ-cyhalothrin both changed average biomass by < 10%).

**Table 1.** Linear mixed effects regression after model selection for *Biomphalaria glabrata* and *Bulinus truncatus* snail biomass in experiment one (*n* = 75). Positive insecticide and fertilizer effects represent top-down and bottom-up effects on snails, respectively. Herbicide effects were neither significant nor consistent with positive bottom-up effects. Infected *Biomphalaria glabrata* biomass did not vary with experimental treatments (see Supplementary Information).

| *Biomphalaria*<br>Biomass | Parameter | Estimate | Std.Error | z-value | p-value |
| --- | --- | --- | --- | --- | --- |
|  | (Intercept) | 1.503 | 0.321 | 4.687 | **<0.001** |
|  | fertilizer | 1.567 | 0.500 | 3.133 | **0.002** |
|  | herbicide | -0.710 | 0.371 | -1.913 | 0.056 |
|  | insecticide | 0.950 | 0.365 | 2.605 | **0.009** |
|  | ZI (Intercept) | -1.609 | 0.438 | -3.678 | **<0.001** |
| *Bulinus*<br>Biomass | Parameter | Estimate | Std.Error | z-value | p-value |
|  | (Intercept) | 0.229 | 0.119 | 1.926 | 0.054 |
| Insecticides | λ-cyhalothrin | -0.227 | 0.203 | -1.119 | 0.263 |
|  | chlorpyrifos | 1.260 | 0.199 | 6.345 | **<0.001** |
|  | malathion | -0.229 | 0.204 | -1.126 | 0.260 |
|  | terbufos | 0.965 | 0.222 | 4.342 | **<0.001** |
|  | esfenvalerate | 0.773 | 0.241 | 3.210 | **0.001** |
|  | permethrin | 1.387 | 0.214 | 6.480 | **<0.001** |
| Herbicides | acetochlor | -0.209 | 0.203 | -1.032 | 0.302 |
|  | alachlor | -0.208 | 0.203 | -1.026 | 0.305 |
|  | atrazine | -0.229 | 0.204 | -1.126 | 0.260 |
|  | metolachlor | -0.229 | 0.204 | -1.126 | 0.260 |
|  | propazine | -0.229 | 0.204 | -1.126 | 0.260 |
|  | simazine | -0.229 | 0.204 | -1.126 | 0.260 |
|  | fertilizer | 0.932 | 0.199 | 4.691 | **<0.001** |
|  | ZI (Intercept) | -2.390 | 0.669 | -3.571 | **<0.001** |

**Figure 1.**
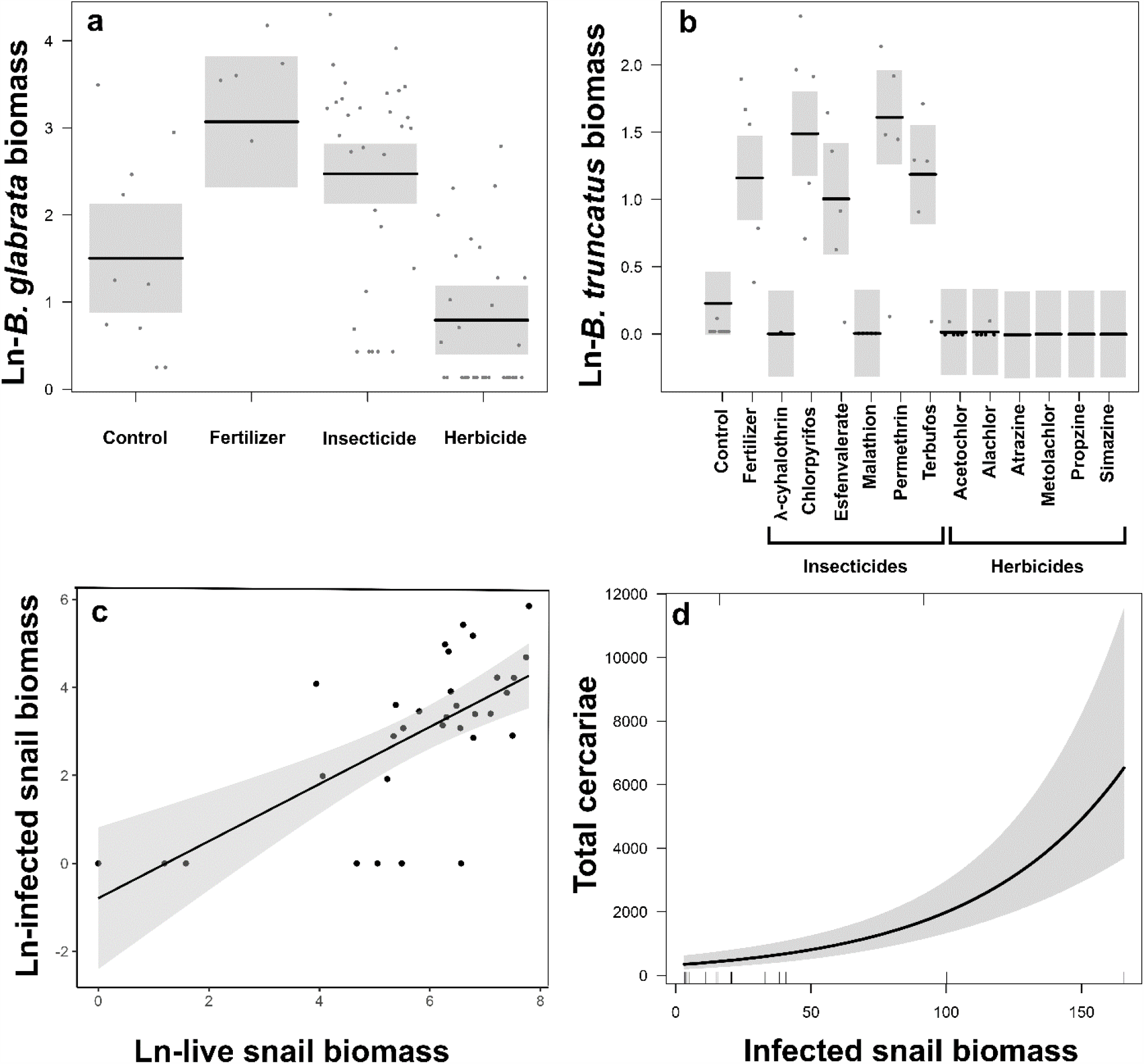
Linear mixed effects model results for natural-log transformed *B. glabrata* (a) and *B. truncatus* (b) biomass in experiment one (*n* = 75) using a Gaussian error distribution, scatterplot of natural-log transformed snail biomass versus infected snail biomass (*n* = 32) in experiment two (c), and a negative binomial model predicting total cercariae using a fixed predictor for infected snail biomass (*n* = 21) in experiment two (d).

Structural equation modeling (SEM) was consistent with the above relationships (Fig. 2). The final SEM was a good fit for *Bu. truncatus* (*CFI* = 0.99, χ^2^ = 15.25, *df* = 26, *p* = 0.95; Table S8) and *Bi. glabrata* (*CFI* = 0.99, χ^2^ = 28.04, *df* = 26, *p* = 0.36; Table S9). The SEM model for *Bu. truncatus* suggested positive and indirect top-down effects of insecticides on snail biomass by lowering 24 hr crayfish survival, which lowered final crayfish abundance (Fig. 2). Fertilizer increased both phytoplankton and periphyton chlorophyll a, the latter of which had positive but non-significant effects upon snail eggs and biomass (Table S8). Instead, significant positive bottom-up effects of fertilizer occurred via increasing submerged vegetation, which increased both snail eggs and biomass (Fig. 2). The sum of standardized coefficients of indirect pathways from fertilizer to snail eggs or biomass was over an order of magnitude greater via submerged vegetation than via periphyton chlorophyll a alone. Submerged vegetation was reduced by crayfish via consumption, which also formed a significant and negative indirect pathway from crayfish predators to snail eggs or biomass that was independent of crayfish predation on snails (Fig. 2). Thus, insecticide-induced reductions in crayfish were positively associated with snails both by reducing predation and increasing snail habitat. The significance and direction of pathways in the SEM model for *Bi. glabrata* were identical to *Bu. truncatus* except that both snail eggs and submerged vegetation pathways were negative for *Bi. glabrata* biomass (Table S9). *Bi. glabrata* biomass at the end of the experiment was also approximately an order of magnitude greater than for *Bu. truncatus* (Table S10).

**Figure 2.**
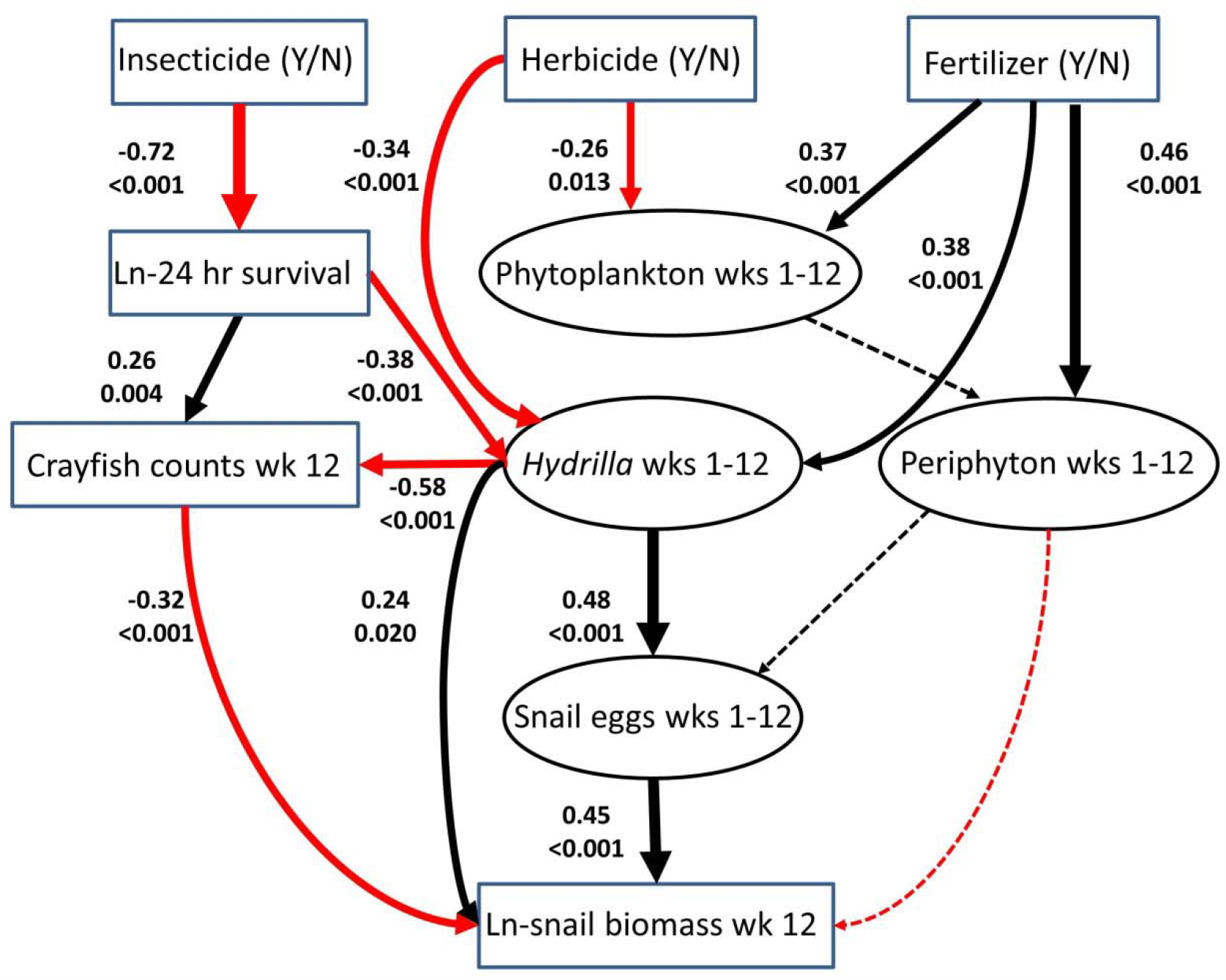
Combined factor and path analysis for *B. truncatus* biomass demonstrating top-down and bottom-up effects of insecticides and fertilizer, respectively. Top-down effects: insecticide exposure reduced final crayfish counts, which directly increased snail biomass by reducing predation, and also indirectly increased snail biomass by increasing snail habitat *H. verticillata* (crayfish ate *H. verticillata*). Bottom-up effects: Fertilizer increased periphyton and *H. verticillata*, which increased snail biomass because they are food and habitat, respectively, for snails. Herbicides reduced *H. verticillata*, snail habitat, and, thus, on average, had net negative effects on snails. P-values for paths in the model are reported below each standardized coefficient. Red arrows denote significant negative paths; black arrows denote significant positive paths; dashed arrows denote non-significant pathways. Boxes represent directly measured variables and circles represent latent variables. Indicator variables for latent variables have been omitted from the figure to reduce visual complexity, but are reported in Table S5. The path model had a good fit to observed data (*CFI* = 0.99, χ^2^ = 26.3, *df* = 26, *p-value* = 0.45).

### Experiment Two

Given that the top-down effects of crayfish were so strong, essentially eliminating juvenile snails, we were concerned that the top-down effects of crayfish were masking any positive effects of herbicides on snail production and schistosomiasis risk. Consequently, we conducted a follow-up experiment where we crossed the same herbicide exposures with the presence and absence of crayfish. However, using five crayfish led to complete depredation of snails in the predator tanks, reducing our number of tanks with live snails in half (*n* = 32). Subsequently, we found that little variance in *Bi. glabrata* biomass was explained by treatment (Table S4), and likelihood ratio tests did not detect significant herbicide effects in our nested models (Table S5). An interaction term between crayfish presence and herbicide type for infected, uninfected, and total *Bi. glabrata* biomass, after model selection (Tables S11-S13), showed no significant interaction between crayfish presence/absence and herbicides (Table 2). Focusing on tanks without crayfish predators, the herbicides atrazine and acetochlor were associated with increases in the average biomass of infected snails (55 %) and total snails (131 %), respectively (Table S13; Fig. 3a,3c). In contrast, the herbicide alachlor was associated with a decrease in uninfected snail biomass (126%) in tanks without predators (Table S14; Fig. 3b). A Pearson’s correlation between snail biomass and infected snail biomass suggested a significant positive association (*rho* = 0.67, *t*_*30*_=4.96, *p* < 0.001; Fig. 1c). Moreover, infected snail biomass was a significant positive predictor of total cercariae in tanks (Fig. 1d).

**Table 2.** Linear mixed effects regression after model selection for infected and uninfected *Biomphalaria glabrata* snail biomass considering an interaction between crayfish predators and herbicides in experiment two (*n* = 64). Herbicide effects did not significantly interact with top-down effects.

| Response | Parameter | Estimate | Std.error | z-value | p-value |
| --- | --- | --- | --- | --- | --- |
| Infected *Biom.* biomass | (Intercept) | 1.53 | 0.11 | 13.4 | <0.01 |
|  | CrayfishYes | -1.45 | 0.16 | -9.01 | <0.01 |
|  | typeherbicide | 0.13 | 0.13 | 1 | 0.32 |
|  | CrayfishYes:typeherbicide | -0.13 | 0.19 | -0.72 | 0.48 |
|  | ZI (Intercept) | -1.68 | 0.42 | -4.01 | <0.01 |
| Uninfected *Biom.* biomass | (Intercept) | 2.67 | 0.17 | 15.69 | <0.01 |
|  | CrayfishYes | -2.59 | 0.24 | -10.7 | <0.01 |
|  | typeherbicide | -0.13 | 0.2 | -0.65 | 0.52 |
|  | CrayfishYes:typeherbicide | 0.13 | 0.28 | 0.46 | 0.65 |
|  | ZI(Intercept) | -3.52 | 1.02 | -3.47 | <0.01 |
| Total *Biom.* biomass | (Intercept) | 3.29 | 0.10 | 31.47 | <0.01 |
|  | CrayfishYes | -1.30 | 0.15 | -8.49 | <0.01 |
|  | typeherbicide | 0.06 | 0.12 | 0.53 | 0.60 |
|  | CrayfishYes:typeherbicide | -0.02 | 0.18 | -0.14 | 0.89 |
|  | ZI(Intercept) | -3.01 | 0.59 | -5.09 | <0.01 |

**Figure 3.**
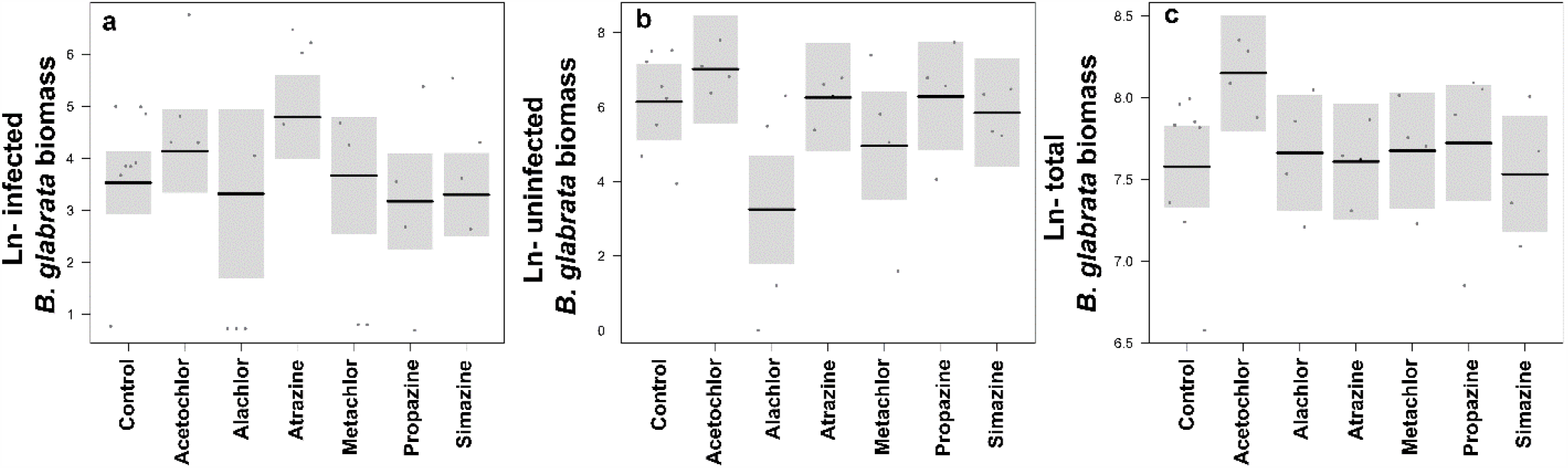
Linear mixed effects model results using a fixed categorical predictor for each herbicide to predict natural-log transformed infected (a), uninfected (b), and total *B. glabrata* biomass (c) in tanks without predators in experiment two (*n* = 32) using a Gaussian error distribution. Only atrazine and acetochlor were positively associated with infected and total *B. glabrata* biomass (*p*<0.05), respectively, whereas alachlor was negatively associated with uninfected *B. glabrata* biomass (*p*<0.05).

## Discussion

Given that recent evidence indicates that increasing agriculture to meet the growing global food demand could simultaneously increase human infectious diseases (Rohr et al., 2019), such as human schistosomiasis, we conducted a study to identify agrochemicals that could sustainably improve food production without increasing schistosomiasis. Expanding upon previous work on predators in isolation (Halstead, Civitello, & Rohr, 2015), our study confirmed that most insecticides pose a high toxicity risk to snail predators at environmental concentrations in simulated aquatic communities. We identified that, within their pesticide class, the organophosphate malathion and the pyrethroid λ-cyhalothrin had the lowest risks of increasing *Bu. truncatus* snails that transmit urinary schistosomiasis in Africa. Bottom-up effects of fertilizer on snail biomass in natural communities occurred as fertilizers increased both snail food (periphyton) and habitat (aquatic vegetation), whereas herbicides decreased snail habitat (experiment one). Thus, fertilizer bottom-up effects are expected to be generally stronger than those for herbicides in natural communities. Positive bottom-up effects on snails for the herbicides atrazine and acetochlor were found in the absence of aquatic macrophytes and snail predators (experiment two), whereas some herbicides, such as alachlor, were negatively associated with snails under the same conditions. We found that snail biomass positively predicted their cercarial production, suggesting that changes in snail biomass associated with agrochemical pollution could have strong potential to influence human exposure to schistosome cercariae. Our findings highlight that maximizing the benefits of agricultural intensification, while minimizing indirect costs on human health, will require identifying bottom-up and top-down effects of agrochemicals within natural aquatic communities where the snails that transmit schistosome parasites are found.

Initial (24hr) mortality of the crayfish *Procambarus alleni* in our mesocosm experiment generally matched expectations from previous lab experiments, with the exception of the toxicity of some pyrethroids being lower in outdoor mesocosms. We confirmed previous findings that malathion is a low risk organophosphate, potentially because of its very high LC_50_ values for *P. alleni* (Halstead et al., 2015). Halstead et al. (2015) reported that all three of the pyrethroids that we tested were highly toxic to *P. alleni*. However, we found that λ-cyhalothrin, unlike other pyrethroids, did not lead to top-down effects on snails in our natural communities when applied at concentrations approximating the 96h LC_50_ for *P. alleni* (Halstead et al., 2015). One potential reason for this result may be that submerged vegetation, such as the *H. verticillata* used in our study, has been linked to reduced toxicity of pH-sensitive insecticides, including malathion and λ-cyhalothrin (Brogan & Relyea, 2014; Leistra et al., 2004). While the organophosphate chlorpyrifos and the pyrethroid esfenvalerate also exhibit pH-sensitive hydrolysis (Hertfordshire, 2013), they were likely applied at sufficiently high concentrations in our study (2x and 10x their respective 96h LC_50_ estimates) to preclude any increased degradation from mitigating acute toxicity to *P. alleni*. Thus, our findings suggest that, of the insecticides we tested, malathion and λ-cyhalothrin should represent the weakest top-down effects on snails in natural communities with aquatic vegetation, and the lowest risk of increasing snail biomass.

Fertilizers can increase submerged vegetation that acts as snail habitat, potentially making fertilizers more likely to have positive bottom-up effects on snails than herbicides and even modifying top-down effects. Unlike other agrochemicals, fertilizers are both commercially available and locally sourced via manure products, leading to their wide application even by rural subsistence farmers. Consistent with previous mesocosm experiments, we found that fertilizers can increase snails by increasing attached algae that snails eat (Halstead et al., 2018). However, we also documented that stronger bottom-up effects of fertilizer exist by increasing the submerged vegetation that serves as snail habitat. By increasing *H. verticillata*, fertilizers indirectly benefited *Bu. truncatus* biomass by increasing both snail reproduction and growth. In contrast, the same associations were negative for *Bi. glabrata*, likely because of resource depletion in some tanks given that *Bi. glabrata* infection rates and biomass were both an order of magnitude greater than for *Bu. truncatus*. In early weeks, algal food resources for snails fueled by fertilizer would be high and available for snail reproduction, growth, and parasite production by infected snails (Civitello, Fatima, Johnson, Nisbet, & Rohr, 2018). As *Bi. glabrata* populations quickly grew, per-capita resources in tanks decline such that snail populations crash as periphyton resources become limiting in microcosms (Civitello et al., 2018; Rohr, Halstead, & Raffel, 2012). Thus, aquatic vegetation likely fueled snail reproduction for both snail species, but *Bi. glabrata* biomass outpaced *Bu. truncatus* biomass, likely increasing the likelihood of population crashes (negative associations) toward the end of our experiment. We know of no available studies documenting that such resource limitations occur in aquatic systems where schistosomiasis is transmitted to humans. In fact, available field studies suggest that aquatic vegetation is highly beneficial to both *Bu. truncatus* and *Bi. glabrata* populations in natural systems and to their parasite production (Haggerty, Bakhoum, Civitello, et al., 2020).

In our first experiment, presence of *H. verticillata* in the tank provided an alternative resource for crayfish (Lodge, Kershner, Aloi, & Covich, 1994), in addition to providing potential refugia and increased resource availability for snails (Bronmark, 1989). Crayfish of the genera *Procambarus* and *Oronectes* have been associated with declines in both submerged macrophytes and snails (Lodge et al., 1994; Rodríguez, Bécares, & Fernández-Aláez, 2003). Reductions in macrophytes by crayfish occur through both consumptive and non-consumptive removal of plant biomass (Lodge et al., 1994; Rodríguez et al., 2003), while reductions in snails are primarily consumptive (Hobbs, Jass, & Huner, 1989). We found that crayfish were negatively associated with *H. verticillata* cover, consistent with crayfish consuming this macrophyte. However, submerged plants actually were associated with reduced final crayfish counts, possibly by providing refugia for snails thus reducing crayfish predation rates on this primary protein source. In addition to providing potential refugia from snail predators, macrophytes provide increased surface area for epiphytic periphyton growth, and grazing of epiphytes by snails can, in turn, increase macrophyte growth (Bronmark, 1989; Lodge et al., 1994; Schmidt, Koehler, & Alfermann, 2011; Underwood, 1991). Nutrients, particularly ammonia, released by snails can also facilitate macrophyte growth (Underwood, 1991). Thus, aquatic vegetation can increase snails by both providing resources to snails and reducing predation rates on snails. In contrast to the potential for fertilizers to foster the mutualism between snails and vegetation, we found that effects of herbicides on snails in natural communities were generally weaker, potentially due to a negative association between herbicides and submerged plants in experiment one.

Herbicides generally had negative effects on snails by reducing aquatic vegetation, with positive bottom-up effects of atrazine and acetochlor only in the absence of aquatic vegetation and snail predators. The positive association that we found between some herbicides, such as atrazine, one of the most common herbicides globally, and snails in our second experiment is consistent with previous studies (Halstead et al., 2018; Rohr et al., 2008). Halstead et al. (2018) actually detected positive atrazine effects while controlling for crayfish density. However, Halstead et al. (2018) did not include submerged aquatic vegetation, which likely increased the chances of detecting bottom-up herbicide effects because it omits the negative herbicide effects on snail habitat that we found in experiment one. Atrazine and other herbicides can increase snails by killing phytoplankton that competes with periphyton algae eaten by snails (Halstead et al., 2018). Instead of such mechanisms leading to consistent positive effects on snails among herbicides, we found positive and negative effects in our study for acetochlor and alachlor, respectively. As algae absorb available nutrients, both periphyton and snails that consume these nutrients, eventually hit a carrying capacity after which their populations can decline or even crash (Rohr et al., 2012). Herbicide effects can be negative overall when considering only data points at the end of the experiment after snail populations exceed carrying capacity and populations subsequently crash (Rohr, 2017). Thus, we think that variation in snail population growth among tanks likely influenced alachlor and acetochlor associations with dead and total snails, respectively, when measured at the end of our second experiment. Nonetheless, our results suggest that fertilizers are far more likely than herbicides to have positive bottom-up effects on snails, and especially in communities where snail predators are present. Our finding that fertilizers can increase snail habitat and resources is consistent with field observations in sub-Saharan Africa that revealed positive associations between fertilizer use and snail habitat, snail abundance, and human schistosomiasis infections (Haggerty, Bakhoum, Chamberlin, et al., 2020).

The importance of other functional groups, such as aquatic vegetation, in mediating the indirect effects of contaminants on snails that transmit the parasites causing schistosomiasis in humans underscores the need to include these relationships when assessing environmental risk of contaminants. This becomes even more critical when assessing the risk of contaminant mixtures. The combined indirect effects of multiple contaminants can either mitigate or exacerbate the predicted effects of a single-contaminant scenario (Halstead et al., 2014). Our study inferences are restricted to observing chemicals individually and we could not simulate all biotic components of the aquatic communities that occur in transmission sites that might influence snail responses to agrochemicals. For example, eutrophication from long-term fertilizer inputs could cause unpredictable changes to snail populations or species richness, potentially eliminating important functional redundancies that can alleviate losses in functionally-important sensitive species (Hooper et al., 2005). Thus, the extent to which changes in biodiversity and abiotic conditions can mediate indirect effects of contaminants (either individually or as mixtures) on *Biomphalaria* and *Bulinus* snails warrants further study. This is especially true given that changes in snail biomass associated with agrochemicals that we found appear very likely to change parasite production. Further, human co-infection by parasites released by *Biomphalaria* and *Bulinus* snails is not uncommon in Africa.

Overall, our findings suggest the need to identify low risk insecticides, such as malathion, that could reduce crop pests without increasing snails. For example, our findings recommend against pesticides that harm crayfish, because these predators reduce transmission directly by consuming snails and indirectly by consuming snail habitat. The more consistent and stronger positive impact of fertilizers on snails than herbicides, by increasing both their food and habitat, highlights that the removal of aquatic vegetation could offer important public health benefits by limiting its bottom-up effects on schistosomiasis risk.

## Methods

### Experimental Design

#### Experiment One

To examine top-down and bottom-up effects in the presence of snail predators, we established experimental freshwater ponds in 75 800L mesocosms filled with 500L of water (mean pH = 9.09) in the summer of 2011 at a facility approximately 20 miles southeast of Tampa, FL, USA. Tap water in mesocosms was allowed to age for 48 hours before being seeded with algae (periphyton and phytoplankton) and zooplankton collected from local ponds. Algal and zooplankton communities were allowed to establish over a four-week period and water was mixed among tanks weekly to attempt to homogenize initial communities before application of agrochemical treatments. Plastic containers with sediment (1 L play sand and 1 L organic topsoil (The Scotts Company, Marysville, OH, USA)) and five rooted shoots of *H. verticillata* were added to each tank on 5 July 2011. Immediately before application of agrochemical treatments on 11 July 2011 (Week 0), we added snails (21 *Bi. glabrata* [NMRI strain] and 12 *Bu. truncatus* [Egyptian strain], provided by NIAID Schistosomiasis Resource Center) and snail predators (2 juvenile *Procambarus alleni* crayfish, 7 *Belostoma flumineum* water bugs, and 3 *Lethocerus* spp. water bugs collected from local ponds) to each tank.

Tanks were randomly assigned to one of fifteen treatments (5 tanks per treatment) including twelve pesticides at their respective EEC (estimated environmental concentration), fertilizer (440 µg/L Phosphorus; 4400 µg/L Nitrogen, observed levels from the field; (Chase, 2003)), solvent control (0.0625 mL/L acetone), and water control in five replicated spatial blocks. Water and solvent controls were used to ensure that any observed effects of pesticides could not be attributed to the presence of solvent. All pesticides were dissolved in acetone and applied at their respective estimated peak environmental concentrations (Tables S1-S2). EEC values were determined using the USEPA’s GENEEC (v2.0, USEPA, Washington, D.C.) software, manufacturers’ label application recommendations, and the physicochemical properties of each pesticide (Table S1-S2).

#### Snail infections

*Schistosoma mansoni* (NMRI strain) and *S. haematobium* (Egyptian strain) eggs were collected from infected Swiss-Webster mice and Siberian hamsters, respectively, housed at the University of South Florida’s College of Medicine Vivarium. Infected rodents were provided by the NIAID Schistosomiasis Resource Center. Five infected mice and hamsters each were euthanatized on dates (30 August 2011, 1 September 2011, and 6 September 2010), and *S. mansoni* and *S. haematobium* eggs were collected from the livers and intestines, respectively. Eggs were isolated from tissue using a handheld immersion blender and collected on a 45-_μ_m sieve. Mature eggs were maintained in 1.4% NaCl to inhibit hatching in a 50 mL centrifuge tube. Eggs were suspended repeatedly using a vortex mixer (Fisher Scientific) and forty 3 mL aliquots were prepared for each schistosome species and added to the tanks within two hours. An additional three aliquots were preserved to quantify the total number of eggs added to each tank. Egg viability was quantified by placing subsamples of the remaining mature eggs in artificial spring water (Cohen, Neimark, & Eveland, 1980) and average egg viability is provided in Supplementary Table S3.

#### Experiment Two

To examine bottom-up effects of the six herbicides in the absence of snail predators, we performed a follow-up experiment in 2015 using the same general experimental setup above in the same location but with a few alterations from 2011: 1) 64 mesocosm tanks were employed with 8 tanks per treatment, including the water and solvent controls, 2) five crayfish predators were randomly assigned to half of the tanks in each treatment, 3) 24 *Bi. glabrata* (NMRI strain) snails were placed into each tank, and 4) tanks lacked *H. verticillata*.

#### Snail infections

To examine the relationship between snail biomass and cercarial production in the 2015 experiment, *Schistosoma mansoni* (NMRI strain) eggs were collected from infected Swiss-Webster mice housed at the University of South Florida’s College of Medicine Vivarium. Infected rodents were provided by the NIAID Schistosomiasis Resource Center. Sixteen infected mice were euthanatized on each date of egg addition to tanks (27 July 2015, 10 August 2015, 24 August 2015 and 8 September 2015), and *S. mansoni* eggs were collected from the livers. Egg isolation, viability testing, and deployment to tanks within 2 h was performed as described above for experiment one. Average egg viability for the four egg harvests in experiment two was 48 percent (Table S3).

Experimental biosafety protocols were approved by USF Biosafety (IBC # 1334) to minimize the risk of human infection via personal protective gear and to minimize snail escape using methods fully described by Halstead et al. (2018). Briefly, personnel wore shoulder length PVC gloves, and mesocosm tanks had an inward-facing top edge, were only filled halfway, and were covered by a heavily weighted shade cloth. The experimental area was enclosed by a chain-link fence with barbed wire, was located > 200m from any waterbodies, and was surrounded by a double silt fence with molluscicide applied between the fencing every two weeks.

#### Data Collection

##### Experiment One

Data collection for algae and snail eggs was made at weeks 1, 2, 4, 6, 8 and 12 of the experiment. Periphyton measurements were recorded from 100 cm^2^ clay tiles suspended vertically 15 cm from the bottom of each tank (approximately 20 cm below the water’s surface), facing south along the northern wall of each tank. Five clay tiles were added to each tank when they were initially filled with water in June 2011 and July 2015. Each visit, a tile was collected and the phytoplankton removed using a scrub brush and wash bottle of de-ionized water to fill a 50mL falcon tube. A separate 50 mL water sample from the center of the tank was collected for phytoplankton abundance. Phytoplankton and periphyton chlorophyll (measured as F_0_), were measured from samples stored in darkness for 1h, using a handheld fluorometer (Z985 Cuvette AquaPen, Qubit Systems Inc., Kingston, Ontario, Canada).

Snail egg masses and hatchling density were estimated using two 15 x 30 cm pieces of Plexiglass placed in each tank; one suspended vertically 10 cm from the bottom of each tank and one resting horizontally along the tank bottom. Visual searches for dead crayfish in tanks with crayfish predators occurred at 24 and 48 hrs after insecticide addition, and upon each snail egg sampling session. At the end of the experiment (week 12), we quantified the number and mass (g) of live snails of each species and live crayfish in predator tanks. All snails and macroinvertebrates were collected and subsequently preserved in 70% ethanol. We determined snail infection under a dissecting microscope by cracking each snail’s shell and inspecting the hepatopancreas and gonad.

##### Experiment Two

Algae, snail eggs, and snail hatchlings were quantified using the same methodology as the first experiment. At the end of the experiment (week 12), we recorded the number and mass of live predators and snails. All snails and macroinvertebrates were collected and subsequently preserved in 70% ethanol. We determined snail infection under a dissecting microscope by cracking each snail’s shell and inspecting the hepatopancreas and gonad. We estimated snail tissue biomass (mg dry weight) using a mass-shell length (mm) regression where mass = 0.0096*L^3^ as fully described by (Civitello et al., 2018). To assess parasite production in the second experiment, we randomly collected 24 snails from each tank between 9 – 12 AM on September 10 and September 24 (days 45 and 59 post initial egg introduction, respectively) using a small net, measured their mass (g), and then shed them individually in artificial spring water in 30 mL beakers in the field for 1 hr. After shedding, snails were returned to their tank of capture. Collected cercariae were stained with Lugol’s solution and counted in the laboratory under a microscope. At the end of both experiments, we added pool shock (71.8% trichloro-s-triazinetrione, Recreational Water Products, Buford, GA, USA) to each tank at 0.15g/L to kill any snails or infective schistosome cercariae (Halstead et al., 2018).

#### Data Analysis

##### Experiment One

We used the *lme4*() package to examine whether pesticide type, class, or chemical identity explained the greatest amount of variation in snail biomass. To do so, we separately predicted infected and uninfected *B. glabrata* and *B. truncatus* biomass using a categorical random term for pesticide, nested within class, nested within type to account for the nested structure of our experimental design (1 | type / class / chemical). These models did not consider the control tanks because they were not hierarchically nested. To determine whether our nested experimental treatments had significant effects on snails, we then performed a likelihood ratio test to compare a full nested model for each snail species to a null (intercept only) linear model that did not include the random effect. If our nested model was significant, we then used the experimental level that explained the most variation in snails as a fixed categorical predictor of either infected or uninfected *B. glabrata* or uninfected *B. truncatus* total snail biomass using separate regressions with a Gaussian error distribution in the *<u>glmmTMB</u>* package (Brooks et al., 2017) in R v4.0 (RCoreTeam, 2013). We did not analyze infected *B. truncatus* biomass due to very low egg viability, and, thus, infection rates for this species (Table S3). For each response, we also fit models including a random term for spatial block and/or a term for zero-inflation. We considered models having water and solvent tanks treated separately or combined as one control group, resulting in 9 possible models for each species. We ranked models using AICc values, and assessed significance of predictors for each response only for the model with the lowest AICc value (Tables S4-S6). We plotted the results of our regression models using the *visreg* package (Rosseel, 2012).

A combined factor and path analysis was used for the first experiment to explore potential causal pathways of agrochemicals on snail egg production and biomass using the *lavaan* package (Rosseel, 2012) in R statistical software (RCoreTeam, 2013). Because our sample size of 75 tanks restricted the number of causal pathways we could infer, we first constructed a latent variable for phytoplankton chlorophyll a (F_0_) in all weeks, periphyton chlorophyll a (F_0_) in all weeks, snail egg counts in all weeks, and *H. verticillata* abundance in all weeks (1-12). For each latent variable, we used modification indices to add any missing covariances until each model fit the observed data (*CFI* > 0.95). The scores for each latent variable model were then extracted using the predict function in lavaan and used for construction of final structural equation models to separately predict *Bi. glabrata* and *Bu. truncatus* biomass (Tables S7-S8). We examined pathways that were previously known for agrochemicals i.e. top down-effects of insecticides on snail predators and bottom-up effects of fertilizer or herbicide on periphyton (Halstead et al., 2018). We also examined d-separate tests and included any potential missing pathways until our model had a good fit to observed data

##### Experiment Two

We performed the same regressions as for experiment one to predict infected, uninfected, and total snail biomass in experiment two using a categorical random term for pesticide, nested within class, and compared it to a null (intercept) model using likelihood ratio tests. To identify which herbicides have strong effects in the absence of predation, we used the fixed categorical predictor for pesticide identity to separately predict natural-log transformed infected, uninfected, and total *B. glabrata* snail biomass, in tanks without crayfish predators, using a Gaussian error distribution in the *<u>glmmTMB</u>* package (Brooks et al., 2017) in R v4.0 (RCoreTeam, 2013). Total biomass included living and dead snails to examine whether herbicides caused increases in snails early in the experiment despite those snails perishing before the end of the experiment.

To test for an interaction between crayfish predators and herbicides in experiment two, we separately predicted log-transformed uninfected, infected, and total *Bi. glabrata* snail biomass, using Gaussian error distributions that considered all possible interactions between a fixed categorical term for crayfish presence and fixed categorical terms for pesticide, class, or type. We also considered a random term for block, a term for zero-inflation, and whether water and solvent tanks should be treated separately or as a single control group, resulting in 20 possible crayfish interaction models. We compared all interaction models to one another, a model using only crayfish presence and a null (intercept only) model using AICc. We also used a Pearsons’ correlation coefficient to assess the relationship between infected and total snail biomass, and performed a negative binomial regression to predict total cercariae counts in tanks using shedding snail biomass.

